# *Drosophila* cap-binding protein eiF4EHP promotes translation via a 3’UTR-dependent mechanism under hypoxia and contributes to fruit fly adaptation to oxygen variations

**DOI:** 10.1101/2022.07.20.500870

**Authors:** Manfei Liang, Clara Hody, Vanessa Yammine, Romuald Soin, Yuqiu Sun, Xing Lin, Xiaoying Tian, Romane Meurs, Camille Perdrau, Nadège Delacourt, Fabienne Andris, Louise Conrard, Véronique Kruys, Cyril Gueydan

## Abstract

Hypoxia induces profound modifications in gene expression program enabling eukaryotic cells to adapt to lowered ATP supply resulting from the blockade of oxidative phosphorylation. One major consequence of oxygen deprivation is the massive repression of protein synthesis, leaving a limited set of mRNAs to be translated. D. *melanogaster* is strongly resistant to oxygen fluctuations, however the mechanisms allowing specific mRNA to be translated in hypoxia are still unknown. Here, we show that *Ldh* mRNA encoding lactate dehydrogenase is highly translated in hypoxia by a mechanism involving its 3’ untranslated region. Furthermore, we identified the cap-binding protein eiF4HP as a main factor involved in 3’UTR-dependent translation under hypoxia. In accordance with this observation, we show that eiF4EHP is necessary for Drosophila development under low oxygen concentrations and contributes to Drosophila mobility after hypoxic challenge. Altogether, our data bring new insight into mechanisms contributing to Drosophila adaptation to oxygen variations.

## Introduction

Adaptation to variations in oxygen concentration is a conserved mechanism in all metazoans as this process is central for the maintenance of cell and tissue homeostasis^1^. At the cellular level, exposure to hypoxia induces a major reprogramming of gene expression to cope with the energy decrease resulting from the inhibition of oxidative phosphorylation^2^. Two major conserved processes contribute to hypoxia-induced gene reprogramming. The first one relies on the transcriptional activation of gene expression mainly controlled by the Hypoxia Inducible Factors (HIFs), leading to the upregulation of a large panel of genes^3^. The second one corresponds to a profound modification of the translation program combining a strong decrease in global protein synthesis to limit energy demand with selective production of proteins involved in the hypoxic response^4^. While the mechanisms underlying HIF-dependent transcriptional activation have been well characterized^5^, the ones governing translation reprogramming are only partially understood^6^.

Mechanisms that allow mRNAs to be translated during hypoxia include cap-independent recruitment of ribosomes through Internal Ribosomal Entry Sites (IRES). However, although hypoxia is frequently described as up-regulating hundreds of genes in various cell types, only a limited number of transcripts are known to carry *bona fide* IRES^7,8,9^. Besides, only few alternative mechanisms mediated by uORF^10^ or by selective partitioning of mRNA to the endoplasmic reticulum (ER) ^11^ have been described to permit selective mRNAs translation at low O_2_.

Messenger RNA translation initiated by recognition of the 5’cap by eIF4E and binding of eIF4G and the DEAD box ATP-dependent RNA-helicase eIF4A is considered as a reference translational model^12^ while alternative translational initiation complexes have also been described^13^. Particularly, eIF4E2 and its metazoan homologs are well described suppressors of cap-dependent mRNA translation but were also shown to mediate mRNA-specific translational activation^14,15^. In the renal human cell line 786-O, Lee’s laboratory has described an RNA-binding complex formed by RBM4, HIF2α, eIF4E2 and eIF4G3 controlling mRNA translation under hypoxic conditions through interactions involving mRNA 3’UTR^14^. This work has opened new perspectives in understanding the control of protein production in hypoxia. However, HIF2α, a central component of this mechanism, is not conserved in invertebrates such as *Drosophila*^16^, where hypoxic translation has been described^17^. Therefore, our understanding of mechanisms promoting hypoxic translation in metazoans remains partial.

Both *Drosophila* larvae and adult flies as well as *Drosophila* S2 cells were shown to be highly resistant to low O_2_ concentrations, making this model organism very attractive to study hypoxic translation^18,19^. We previously identified *Ldh* encoding lactate dehydrogenase as a highly regulated gene upon oxygen variations in S2 cells^19^. In the present study, we characterized the mechanism promoting translation of *Ldh* mRNA under hypoxic conditions in *Drosophila*. We show that *Ldh* mRNA is efficiently translated under hypoxia despite a strong blockade of protein synthesis. Using a reporter gene approach, we demonstrate that *Ldh* mRNA 3’UTR is sufficient to promote reporter mRNA association to polysomes and translation under hypoxia. Furthermore, we identified the cap-binding translation initiation factor eiF4EHP, the homolog of human eIF4E2 as necessary for the 3’ UTR-dependent translation of *Ldh* mRNA under hypoxia. In accordance with these observations, we found a significant fraction of eiF4EHP to be associated to polysomes and to be excluded from stress granules and P-bodies where most mRNAs are concentrated upon hypoxia. Finally, we demonstrated that eiF4EHP is necessary for *Drosophila* development under hypoxic conditions and strongly influences adult fly mobility upon oxygen recovery.

Altogether, our data reveal a new mechanism promoting mRNA alternative translation upon general inhibition of protein synthesis in *Drosophila* and identify eiF4EHP as a major component of the alternative translation machinery which contributes to *Drosophila* adaptation to oxygen variations.

## Results

### *Ldh* mRNA is efficiently translated under hypoxia by a 3’UTR-dependent mechanism

Increased LDH production is a common feature of oxygen-deficient animal cells. LDH catalyses the reduction of pyruvate to lactate necessary to NAD^+^ regeneration and plays a central role in shifting the cellular metabolism from oxidative phosphorylation to lactic glycolysis upon reduction of oxygen availability^20^.

In D. *melanogaster, Ldh* (also referred as *ImpL3*) is the unique gene coding for the Lactate Dehydrogenase enzyme. We analyzed the production of LDH in *Drosophila* S2 cells after exposure to variable concentrations of atmospheric oxygen. While hypoxia induces a strong reduction of protein synthesis (Fig. 1 A,B), it markedly stimulates LDH production (Fig. 1C). Polysome profiling analysis confirmed that *Ldh* mRNA is largely found in polysomal fractions under hypoxic conditions in contrast to mRNA encoding ribosomal protein RPL32 whose polysomal abundance is decreased in hypoxia (Fig. 1D,E). Therefore, *Ldh* mRNA is efficiently translated in S2 cells exposed to a hypoxic environment while global protein synthesis is strongly repressed under these conditions.

**Figure 1:**
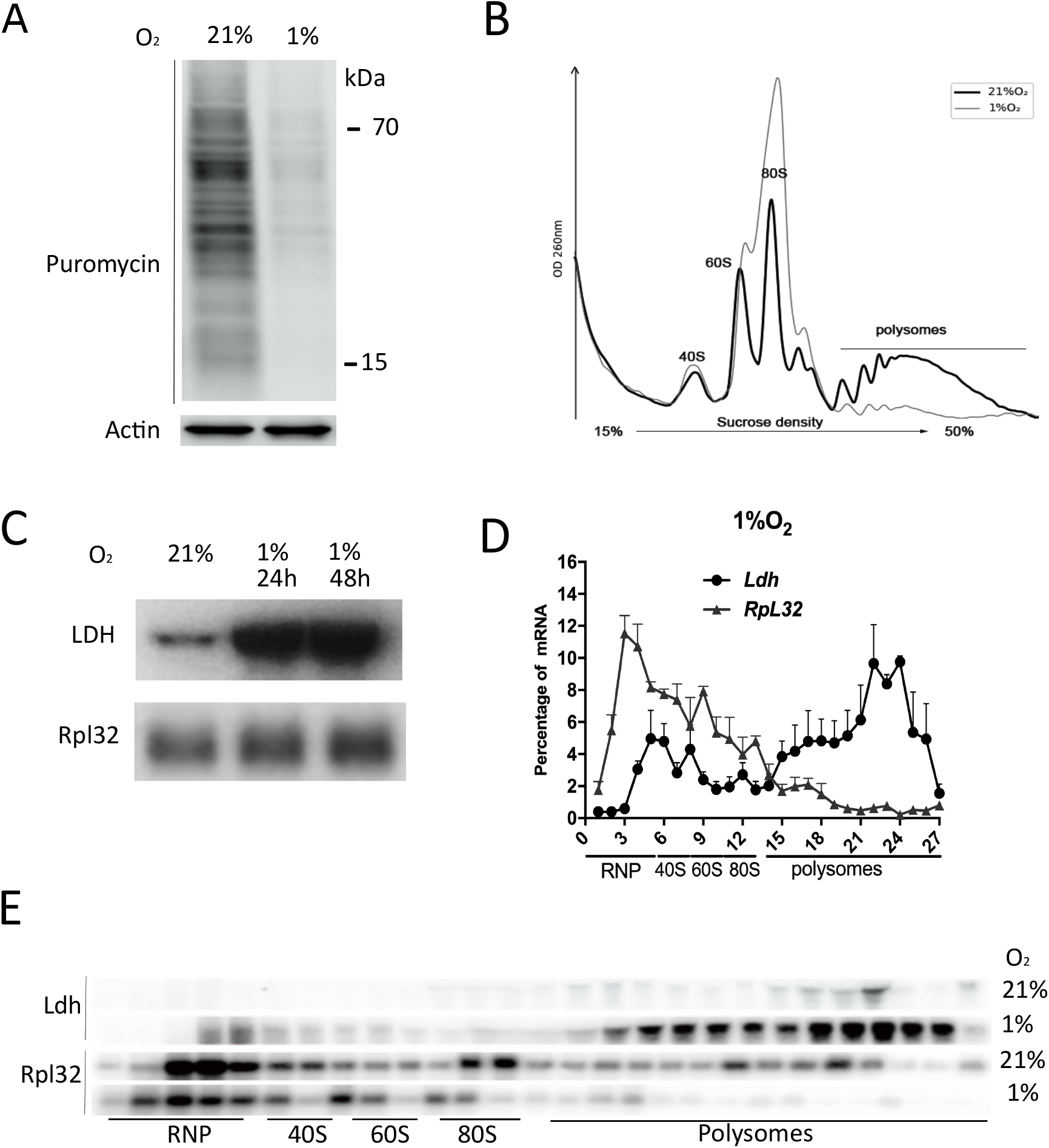
Ldh mRNA is efficiently translated under hypoxia. S2 cells were incubated at 21% or 1% O_2_ concentrations for 24h hours. A) Puromycin was added 20 min. before cell harvesting. Total puromycin incorporation and Actin level were revealed by western blot. (B) Polysome profiles (OD 260 nm) from normoxic (black line) or hypoxic (gray line) cell extracts. Positions of 40S, 60S, 80S and polysomal peaks are indicated. (C) LDH and RpL32 protein levels were detected by western blot in total cell extracts from S2 cells exposed to normoxic conditions or to 1% O_2_ environment for 24 and 48 hours. (D) *Ldh* and *Rpl32* mRNA levels in fractions from polysome profiling experiments were analysed by northern blotting using ^32^P-labeled probes. Experiment was repeated and quantified, data are shown as mean (n=3) ± SEM. (E) One representative experiment performed as in (D) is shown.

We previously demonstrated that *Ldh* mRNA 3’UTR controls the stability of this messenger in normoxic conditions^19^. Reporter gene analysis was performed to evaluate the impact of 3’UTR region in controlling *Ldh* expression in hypoxic S2 cells. Constructs containing the Firefly luciferase (Fluc) coding sequence flanked by different 3’UTR were co-transfected in S2 cells with a Renilla luciferase (RLuc) normalizing reporter plasmid. Quantification of relative luciferase activities after exposure to 1% O_2_ for 24h reveals a significant reduction in Fluc production from reporter mRNA containing the SV40 3’UTR as compared to *Ldh* 3’UTR (Fig. 2A). Reporter mRNAs accumulate at comparable levels in these cells (Fig. 2B), suggesting a difference in translational efficiency. Polysome profiling experiments on stably transfected cells confirmed an efficient accumulation of Fluc mRNA containing the *Ldh* 3’UTR in polysomes of hypoxic cells. In comparison, reporter mRNA containing SV40 3’UTR sediments in polysomal fractions from cells cultivated in 21% O_2_ environment but is largely absent from these fractions upon cell exposure to 1% O_2_ (Fig. 2 C,D). These results indicate that the 3’UTR of *Ldh* mRNA counteracts the general translational blockade imposed by hypoxic stress and is sufficient to maintain an efficient translation rate of *Ldh* mRNA under low oxygen concentration.

**Figure 2:**
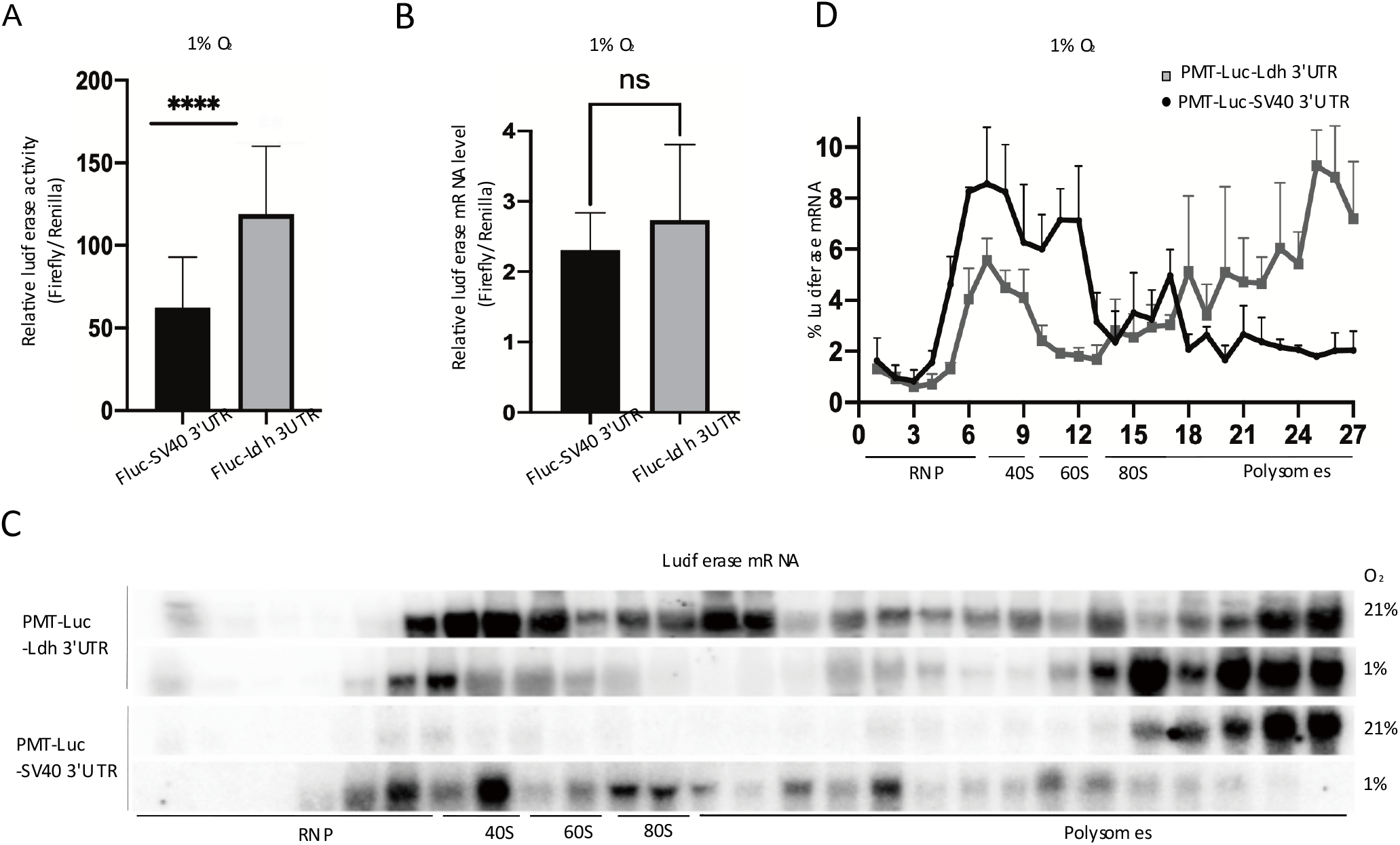
Ldh mRNA 3’UTR enhances translation under hypoxic conditions. S2 cells were transiently transfected with the indicated constructs and exposed to 1% O_2_ for 24h. (A) Firefly (Fluc) and Renilla (Rluc) luciferase activities were measured in total cell extracts by dual luciferase assay and the ratio of both measures was calculated for each transfection. (B) Relative FLuc and RLuc mRNA levels were measured by RT-qPCR. Values represent mean ± SEM, n=3, P-value of one-way ANOVA, ***-P ≤ 0.001, ns-P>0.05, (C) Indicated stably transfected cell lines were exposed to 21% or 1% O_2_ for 24h before fractionation of cell extracts by ultracentrifugation on linear sucrose gradients (15%-50%). The distribution of FLuc mRNA was measured by northern blot in each cell types. (D) Signals from 3 independent experiments performed in 1% O_2_ conditions were quantified and relative signal in each fraction is shown as mean ± SEM.

### eiF4EHP promotes the 3’UTR-dependent translation of *Ldh* mRNA in hypoxia

The *Drosophila* eiF4E1-related protein eiF4EHP and its mammalian homolog eiF4E2 were primarily described as translational repressors of several mRNAs. However, eiF4E2 has also been reported to specifically activate mRNA translation in different cellular contexts including hypoxia (reviewed in^21^).

Interestingly, hypoxic treatment induces an increase of eIF4EHP detected in both head (Fig.3A) and entire body extract (not shown) of adult *Drosophila* and in extracts of S2 cells (Fig. 3B). We therefore investigated the impact of eiF4EHP on translation in hypoxic conditions.

**Figure 3:**
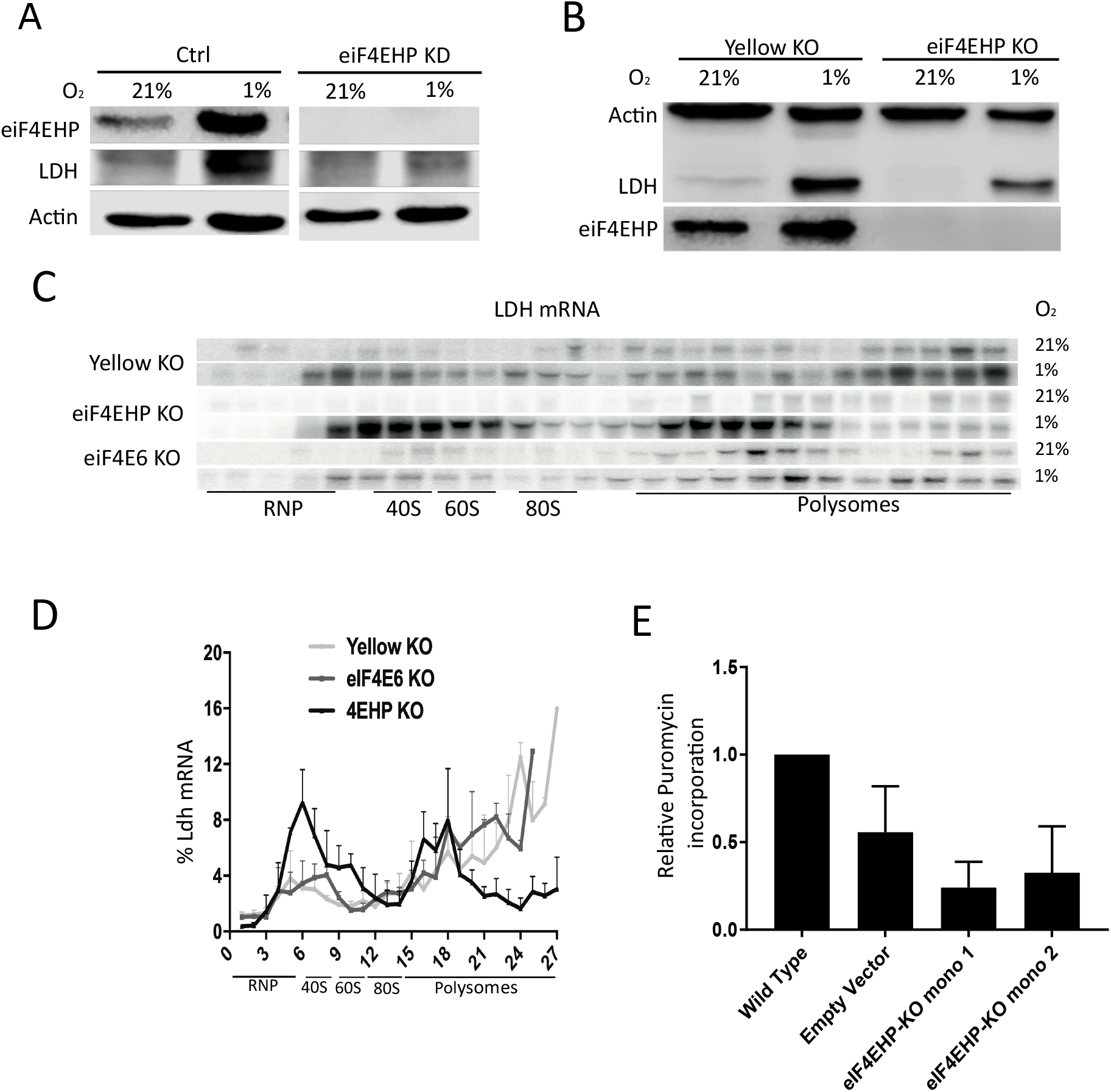
Translational control of LDH production by eiF4EHP in adult flies and S2 cells under hypoxia. (A) Control (Tub>+) or *eiF4EHP* KD (Tub>4HP_RNAi) flies were maintained in either normoxia or hypoxia (1% O_2_, 24h). Protein extracts were prepared from whole flies and eiF4EHP, LDH and Actin were detected by western blot. (B) S2 cells lines inactivated for *Yellow* or *eiF4EHP* genes were established by a CRISPR-Cas9 strategy as described in the methods section. After exposure to normoxic or hypoxic conditions, cells were lysed and Actin, LDH and eiF4EHP protein levels were detected by western blot. (C) Cell extracts from *Yellow, eiF4EHP* or *eiF4E6* KO cells incubated at 21 or 1% O_2_ were separated by centrifugation on linear sucrose gradients and *Ldh* mRNA was detected by northern blot in the different fractions. (D) The mean relative value (± SEM) of *Ldh* mRNA in fractions from 3 independent experiments performed at 1% O_2_ as in (C) is shown. (E) Indicated S2 cell lines were cultivated at 1% O_2_ for 24h and labelled with puromycin (5μg/ml) for 20 min. before harvesting. Puromycin signal in total cell extracts was quantified by western blot and expressed relative to the level detected in wild-type cells.

*Drosophila eiF4EHP* null mutants are not viable^22^. We therefore evaluated the impact of eIF4EHP depletion on LDH production under hypoxic stress by RNAi-mediated knock-down. Adult flies ubiquitously expressing an *eiF4EHP-RNAi* controlled either by a Tub-Gal4 (Tub>4HP_RNAi) or Act5c-Gal4 (Act5c>4HP_RNAi) driver were exposed to normoxic (21% O_2_) or hypoxic (1% O_2_) conditions and LDH was detected by western blot. Control flies (Tub>+; Act5C>+) were exposed to the same conditions. As shown in figure 3A, LDH accumulates after 24h exposure to 1% O_2_ in control 4HP-RNAi flies in the absence of Gal4 driver or in flies expressing a mCherry RNAi controlled by a Tub-Gal4 driver (Tub>mCherry) (not shown). In contrast, LDH level is specifically and strongly reduced in *eIF4EHP RNAi*-expressing flies. LDH production was also reduced in eiF4EHP-deficient S2 cells (Fig. 3B). In accordance with this latter result, the association of *Ldh* mRNA to polysomes is markedly reduced in *eiF4EHP* mutant S2 cells but not in *Yellow* and *eiF4E6* KO cells (Fig. 3C,D). Finally, comparison of puromycin incorporation between *eiF4EHP* KO cell lines and control or wt cells shows that protein synthesis in hypoxia is reduced in the absence of eiF4EHP (Fig. 3E).

As 3’UTR is required for efficient translation of *Ldh* mRNA in hypoxia, we evaluated whether this 3’UTR-mediated mechanism was dependent on eiF4EHP. As shown in figure 4A and B, the association of a Fluc mRNA containing *Ldh* 3’UTR to polysomes observed under hypoxia is disrupted in *eiF4EHP*-deficient cells where most of the reporter mRNA is shifted in non-polysomal fractions. This observation points to a positive effect of eIF4EHP on 3’UTR-dependent translation under hypoxia. To confirm that eIF4EHP specifically affects the expression of the luciferase reporter gene containing *Ldh* 3’UTR, we overexpressed in WT or *eIF4EHP-KO* S2 cells a construct expressing either the D. *persimilis* eIF4EHP protein or GFP as control. The low degree of conservation between D. *melanogaster* and D. *persimilis eiF4EHP* coding sequences in the region targeted by the sgRNA used to establish the KO cells allows the efficient production of D. *persimilis* eiF4EHP in both KO and control cells (Fig. 4C). Cells were transiently co-transfected with a Fluc-Ldh 3’UTR reporter plasmid and a Rluc normalizing plasmid and were further incubated in hypoxia for 24 hours. We observed an increased Fluc activity in presence of D. *persimilis* eIF4EHP as compared to GFP both in *eIF4EHP-KO* and WT S2 cells (Fig. 4D). Similar levels of Fluc mRNA were detected in cells expressing GFP or D. *persimilis* eiF4EHP (Fig. 4E), thereby confirming a translation activating effect of D. *Persimilis* eiF4EHP on Fluc expression. Taken together, these data demonstrate that translation of *Ldh* mRNA in hypoxia is controlled by its 3’UTR and requires eiF4EHP.

**Figure 4:**
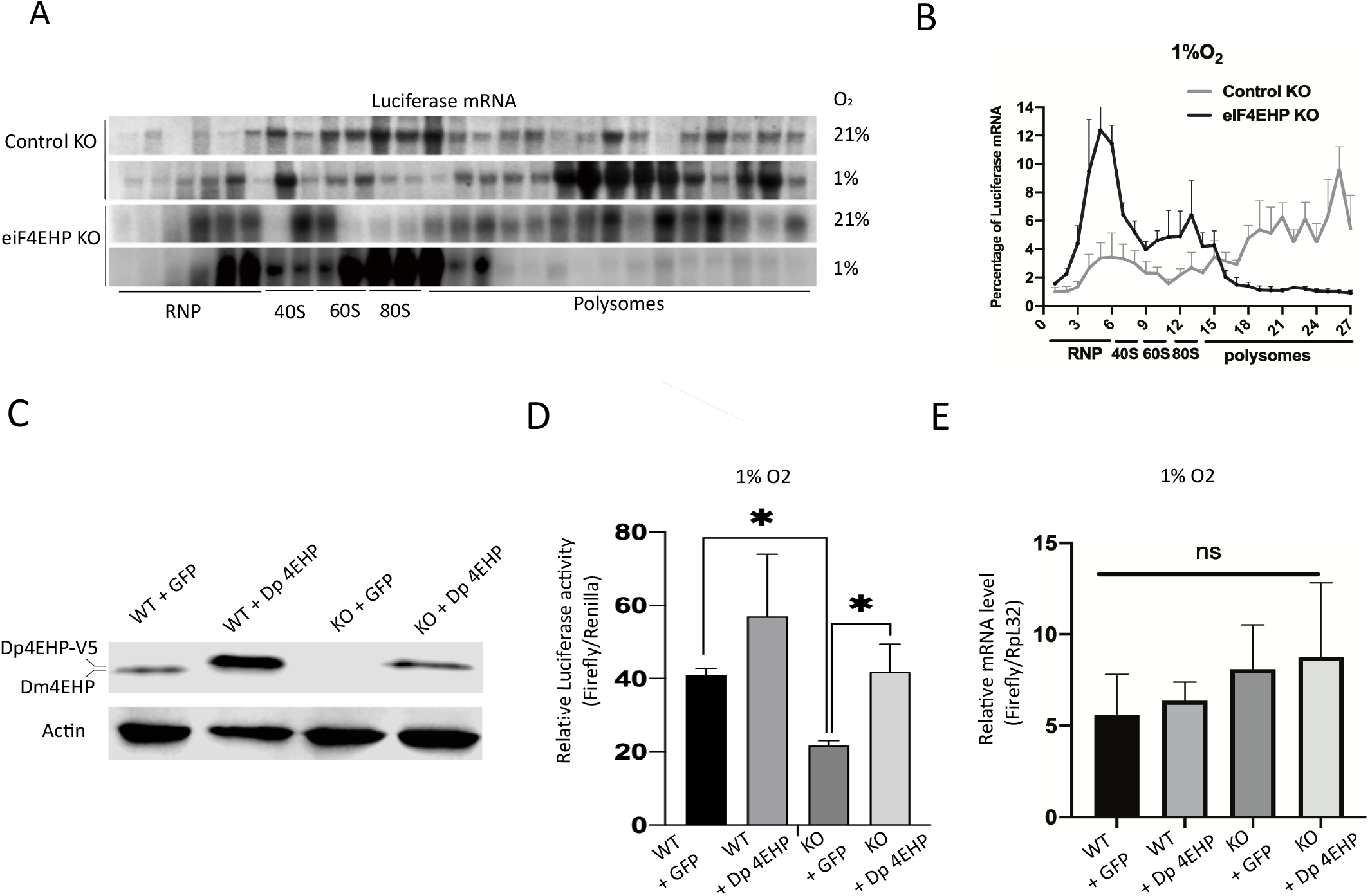
3’UTR-dependent translational control by eiF4EHP in hypoxia. (A) Polysome profiles of FLuc-Ldh 3’UTR reporter mRNA in normoxic and hypoxic *eiF4EHP* KO and control (*Yellow*) KO S2 cells. (B) Quantification of luciferase mRNA levels from 3 independent experiments performed at 1% O_2_ as in (A). (C) WT and *eiF4EHP* KO cells were co-transfected with plasmids encoding V5-tagged D. *persimilis* eiF4EHP (Dp 4EHP) or GFP, along with FLuc-Ldh3’UTR and RLuc reporters. eiF4EHP proteins were detected by western blot. 24h after transfection, cells were cultivated in hypoxia (1%O_2_) for an additional 24h before measurement of luciferase activity by dual luciferase reporter assay (D) and luciferase mRNA levels by qPCR (E). Values represent mean ± SEM, n=3, P-value of one-way ANOVA, *-P ≤ 0.05, ns-P>0.05.

### eIF4EHP is enriched in cytoplasmic foci distinct from SG and P bodies in hypoxia

Hypoxic stress is an inducer of stress granule (SG) assembly in both human and *Drosophila* cells^23,24^. Upon stress, polyadenylated mRNAs accumulate in SG and are mostly excluded from the translational pool ^25,26^. Several members of the eiF4E family were previously shown to relocalize to SG upon cellular stress. However, eiF4E2, the mammalian homolog of eiF4EHP, displays a variable subcellular localization depending on cellular stress types^27^. We investigated the subcellular localization of eiF4EHP and mRNA in response to hypoxic stress. Upon hypoxia, polyadenylated mRNAs massively relocalize to cytoplasmic structures compatible with the previously described concentration of mRNA in SG (Figure 5 A, right panel). As expected, *Ldh* mRNA detection is increased after hypoxic treatment, but the cytoplasmic distribution of *Ldh* mRNA remains more diffuse as compared to the bulk of polyadenylated mRNAs (Fig. 5A, left and middle panels).

**Figure 5:**
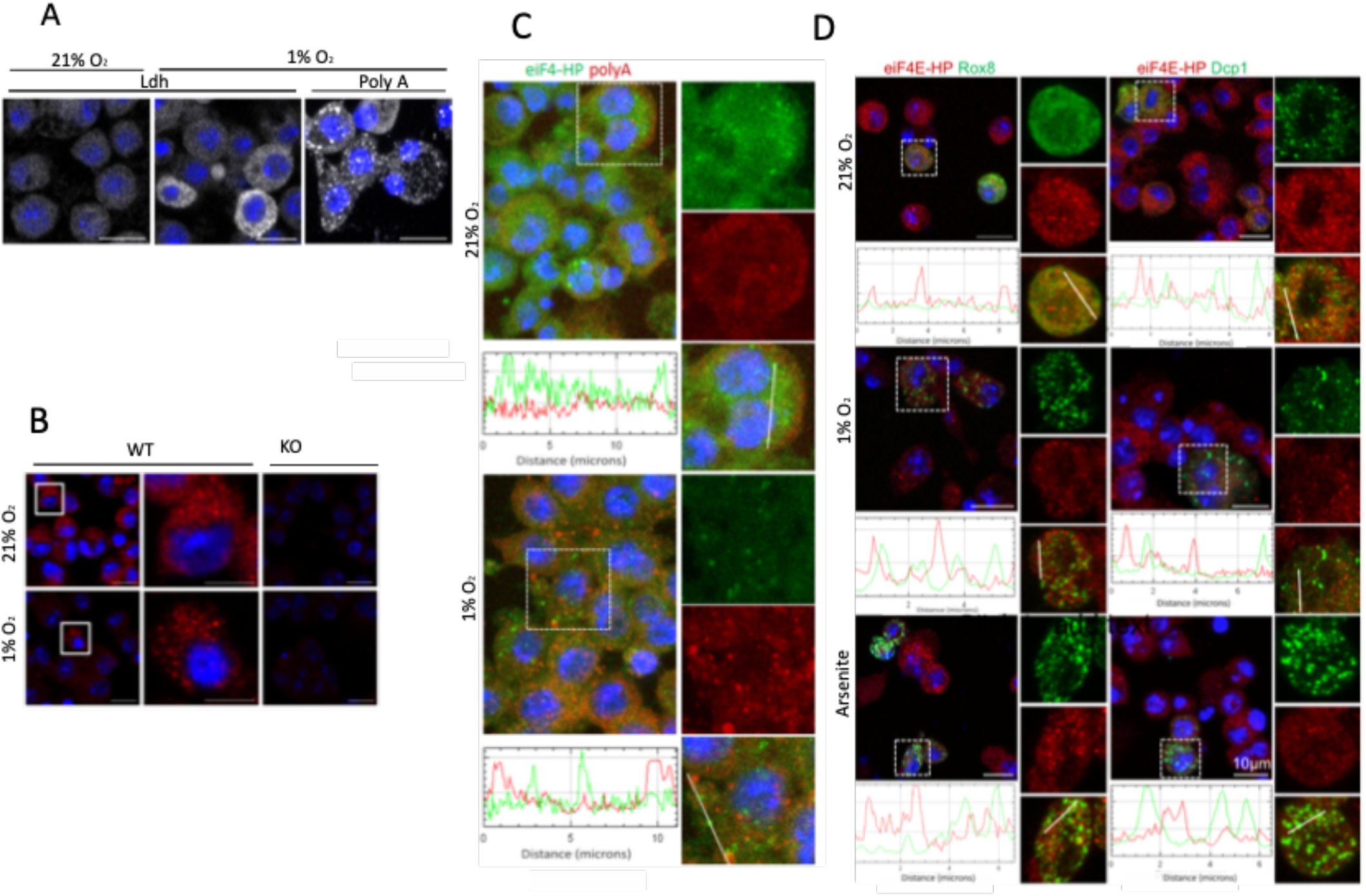
eIF4EHP accumulates in cytoplasmic granules distinct from p-Bodies and stress granules in hypoxia. (A) WT S2 cells were cultivated in normoxia (21% O_2_) or hypoxia (1%, 24h) before analysis by single molecule FISH to detect *Ldh* mRNA (left and middle panels) or Oligo dT FISH to detect polyadenylated mRNAs (right panel). (B) Immunofluorescent detection of ei4EHP in WT or *eiF4EHP* KO cells cultivated at 21% or 1% O_2_ (24h) using anti-eiF4EHP monoclonal antibody (see methods). (C) Combined detection of eiF4EHP by immunofluorescence (green) and polyA mRNA by oligo-dT FISH (red) in normoxic and hypoxic WT S2 cells. (D) Combined detection of GFP-Rox8 or GFP-Dcp1 (green) and endogenous eiF4EHP (red) in transfected S2 cells cultivated in normoxia (21% O_2_), in hypoxia (1% O_2_ for 24h) or after oxidative stress (250μM arsenite for 1h). Representative images of confocal imaging (Z-stack maximum intensity projection) and zooms are shown. In all experiments, nuclei were counterstained with DAPI (blue). Red and green signal intensities were quantified with the Zen software (Zeiss).

In normoxia, eiF4HP has a heterogeneous distribution in the cytoplasm of S2 cells that is further enhanced after exposure to hypoxia (Fig. 5 B,C). The combined detection of eIF4EHP and polyA^+^ RNA confirmed that hypoxic treatment increased the granular distribution of eiF4EHP and mRNAs in the cytoplasmic compartment. However, co-localization analyzes revealed that these markers remain separately distributed under hypoxia (Fig. 5C). To further characterize the nature of the eiF4EHP-containing foci, we performed co-localization experiments with ectopically expressed GFP-Rox8 and GFP-Dcp1 fusion proteins which are *bona fide* markers of stress granules and PB, respectively^28^. As shown in Figure 5D (left panels), GFP-Rox8 is homogeneously distributed in the cytoplasm of S2 cells in normoxia and accumulates in cytoplasmic granules upon hypoxic or arsenite treatments while eIF4EHP remains mostly excluded from these structures. Similarly, P-bodies identified by GFP-Dcp1 labelling, displayed very partial co-localization with eIF4EHP in all tested conditions (Fig. 5D, right panels). Altogether, these data indicate that in hypoxia, eIF4EHP does not colocalize with the bulk of mRNAs and accumulates in cytoplasmic foci distinct from SG and P-bodies.

### eIF4EHP is associated to polysomes and binds to Ldh mRNA under hypoxia

To further investigate the function of eiF4EHP in translation under hypoxic conditions, we analyzed its distribution into polysomes by centrifugation on sucrose gradient. As shown in Figure 6A and B (left panels), a large proportion of eif4EHP co-sediments with light fractions corresponding to ribonucleoprotein particles (RNP), ribosomal subunits (40S/60S) and assembled ribosomes (80S) in extracts from S2 cells and adult fly heads in normoxia. However, under hypoxia, an increased proportion of eiF4HP is found in polysomes in contrast to ribosomal RpS23 protein which shifts from polysomes to lighter fractions (right panels). These data suggest that eiF4EHP is recruited to the translating machinery under hypoxic conditions. To further demonstrate the direct role of eiF4HP in translation under hypoxia, we tested its association with *Ldh* mRNA by crosslinking immunoprecipitation (CLIP). Protein extracts from hypoxia-treated WT and *eIF4EHP-KO* cells were immunoprecipitated with agarose anti-eiF4EHP-coated beads after UV crosslinking and the presence of *Ldh* and *Rpl32* (control) mRNA was analyzed in inputs and eluted fractions by RT-qPCR. As shown in Figure 6C, *Ldh* mRNA was enriched in the eluate from WT crosslinked extract as compared to corresponding fractions of *eiF4EHP-KO* extracts. In contrast, no enrichment of *Rpl32* mRNA was detected in the eluate from immunoprecipitated extracts, indicating that eIF4HP specifically interacts with *Ldh* mRNA in WT cells under hypoxia.

**Figure 6:**
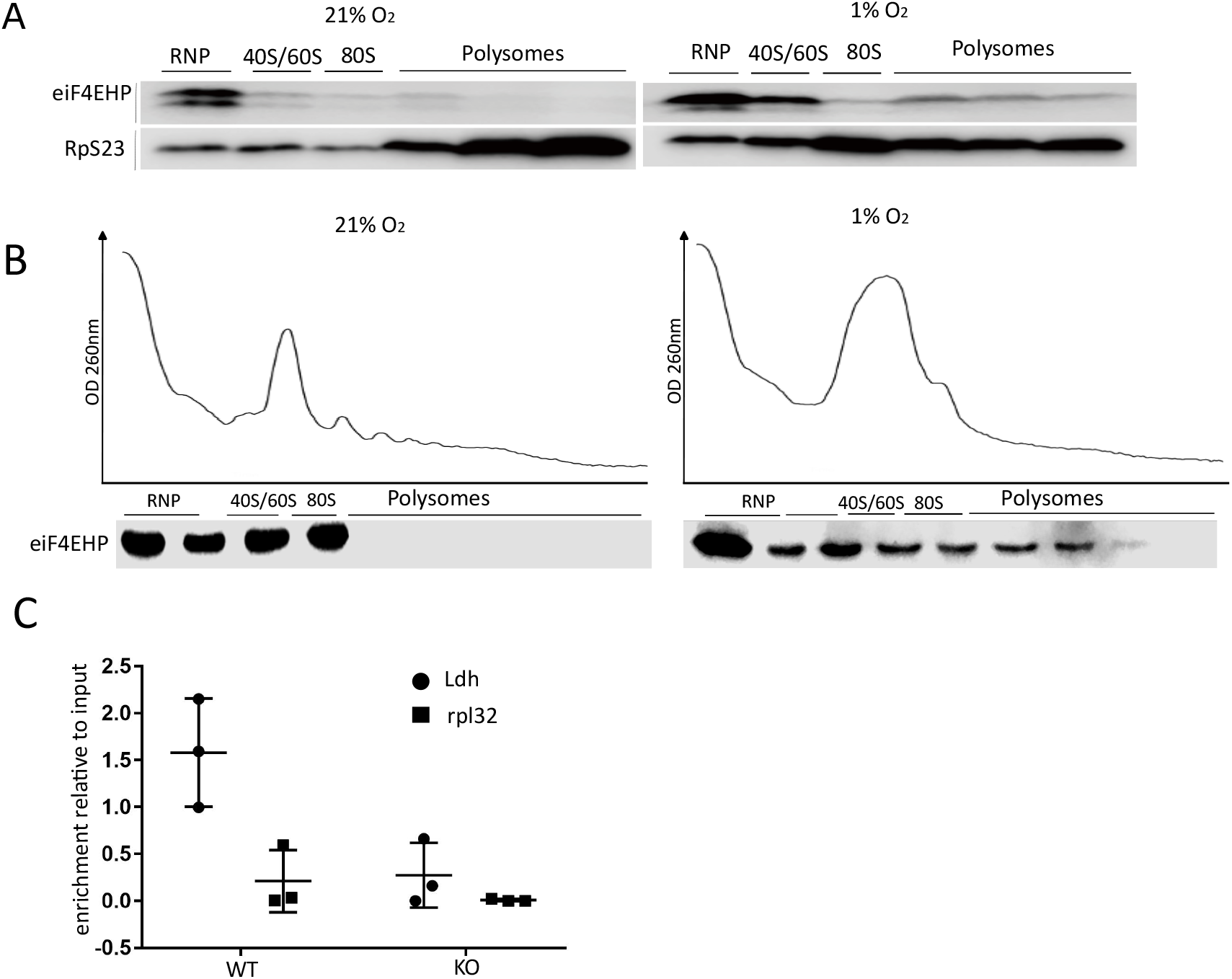
eiF4EHP is recruited to polysomes and interacts with *Ldh* mRNA in hypoxia. (A) Cell extracts from normoxic or hypoxic S2 cells were ultracentrifuged as described in fig.1B, protein extracts from pooled fractions corresponding to the same ribosomal content were analyzed by western blot with anti-eiF4EHP and RpS23 antibodies. (B) Adult Canton S flies were kept at 21% or 1% O_2_ for 24h. Total protein extracts were prepared as described in the method section and fractionated by ultracentrifugation on sucrose gradients (15-50%). OD at 260 nm was measured in line during fractionation (upper panels). EiF4EHP was detected by western blot in representative fractions of the gradients. (C) Cross-Linking and Immunoprecipitation experiment (CLIP) was performed on cell extracts from normoxic and hypoxic (1% O_2_, 24h) UV irradiated WT or *eiF4EHP* KO S2 cells. eiF4EHP was immunoprecipitated with eiF4EHP antibody-coated protein G sepharose beads before *Ldh* and *Rpl32* mRNA detection by RT-qPCR in total extract (input) or immunoprecipitation eluate. mRNA enrichment levels in eluate as compared to input from 3 independent experiments are shown.

Taken together, these results indicate that ei4EHP is involved in the direct recruitment of specific mRNA to the translation machinery under conditions where the global translation activity is strongly reduced.

### eIF4EHP is required for development under hypoxic environment and for recovery from hypoxic exposure in D. *melanogaster*

D. *melanogaster* has a high tolerance to oxygen deprivation both during developmental and adult stages^29,30^. To evaluate the importance of eIF4EHP on hypoxia tolerance, flies carrying *UAS-eiF4EHP* or *UAS-mCherry* RNAi cassettes were crossed with Tub-Gal4/Tm3 or Actin5C-Gal4/Tm6b flies. 24h after egg laying, embryos were kept in normoxia or exposed to 6% O_2_ environment until hatching. *eIF4EHP* knock-down or WT individuals were sorted based on the expression of balancer phenotypical markers. In normoxia, the number of *eiF4EHP* KD individuals exceeded the number of control individuals bearing a balancer chromosome. This difference could be explained by the general reduced transmissibility of balancer over WT chromosomes^31^. As expected, under hypoxia, a global reduction of birth rate is observed independently of the genotype. However, the number of *ei4EHP* KD individuals was strongly reduced as compared to controls with a more pronounced effect upon maternal transmission of the *ei4EHP*-RNAi allele for which no *eIF4EHP* KD individuals were detected (Fig. 7A).

**Figure 7:**
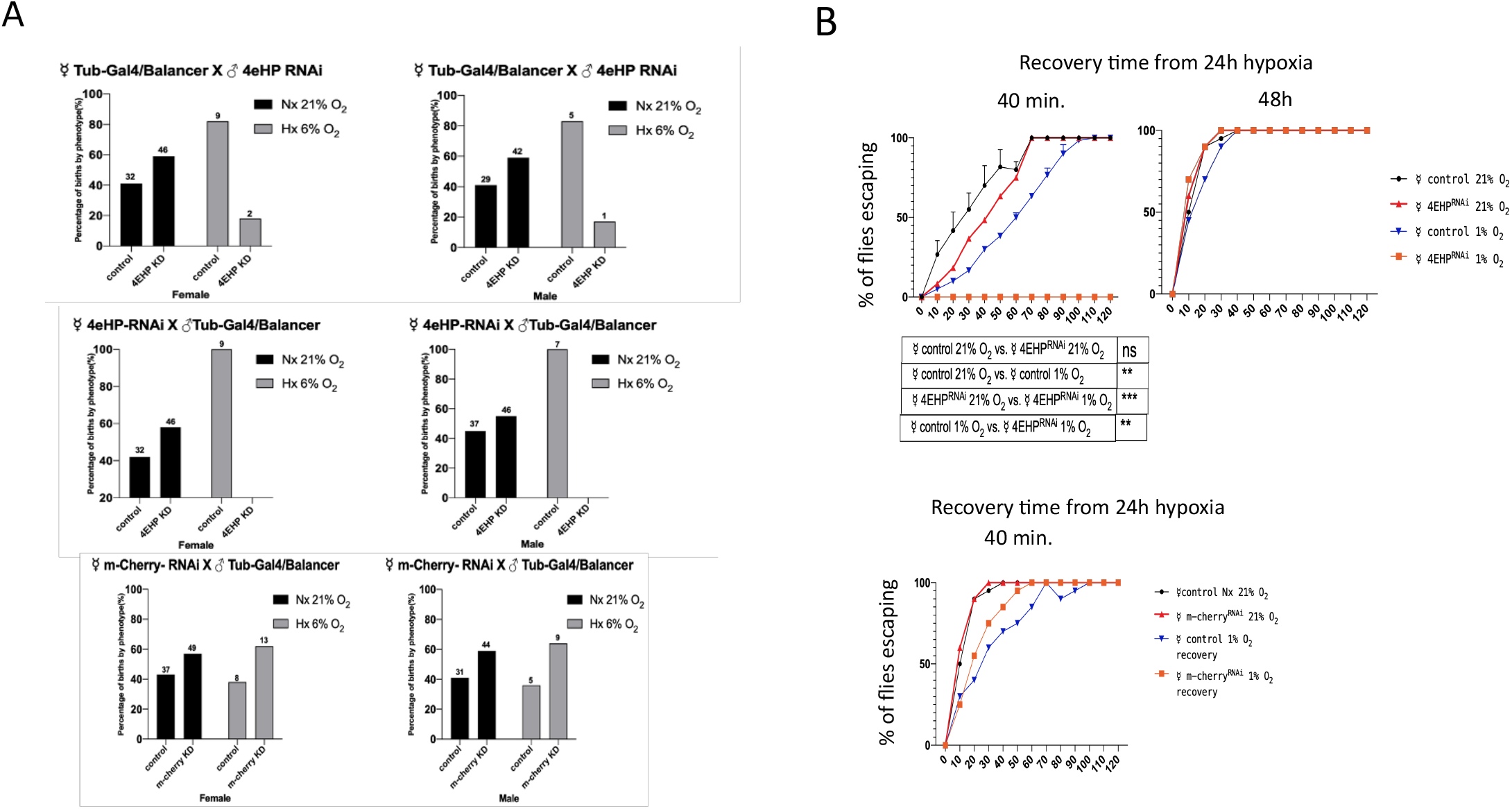
Role of ei4EHP in development under hypoxia and mobility recovery after hypoxia exposure in D. melanogaster. (A) Female or male flies with chromosomal insertion of an UAS_eiF4EHP (BDSC 36876) or UAS_ mCherry (BDSC 35785) RNAi construct were crossed with male or female Tub>Gal4/Tm3 (BDSC 5138) driver flies. 24h after mating, embryos were transferred to 6% O_2_ or kept in normoxia until hatching. Tub-Gal4/UAS-eiF4EHP and Tm3/UAS-eiF4EHP F1 individuals were counted based on the presence (control) or absence (KD) of the Tm3 SB marker and results were expressed as % of total births. (B) Mobility of age-matched Tub-Gal4/UAS-eiF4EHP (4EHP^RNAi^), TM3/UAS-eiF4EHP (control) (upper panels) or TubEHP-Gal4/mCherry_RNAi (m-cherry^RNAi^) (lower panel) females was assessed by negative geotaxis assay in normoxic conditions (21% O_2_) or 40 min. and 48h after an initial exposure to 1% O_2_ for 24h. Flies crossing the climbing limit were counted as “escaping” and expressed as % of total flies. Each assay was performed with 50 flies.

Exposure to hypoxic conditions alters a large panel of physiological parameters in *Drosophila*^32^. Adult flies can survive to strong but transient reduction of O_2_ concentration but show impaired mobility during the recovery period^33^. We tested the effect of eiF4EHP depletion on this recovery of mobility after hypoxia. Adult *eiF4EHP-KD* or control flies were exposed to a 1% O_2_ environment for 24 hours before returning to normoxic conditions. The mobility capacity of flies was assessed by a negative geotaxis test 40 minutes after return to normoxia. All fly strains show comparable mobility when kept in normoxic conditions (Fig. 7B, left panel). *eIF4EHP* KD flies show a strong reduction of mobility during the recovery period with no fly reaching the preset climbing limit during the time of the assay (Fig. 7B, left panel). This defect in mobility was totally recovered 48h after the hypoxic period (Fig. 7B, right panel). Control flies which do not express *eIF4EHP* RNAi have a reduced mobility after 40 minutes of recovery in normoxia but not to a comparable extent to that observed for *eiF4EHP* KD flies. Finally, expression of a control *mCherry* RNAi does not alter flies’ recovery from hypoxic exposure as illustrated in Figure 7B (lower panel). Altogether, these data show that eIF4EHP is important for the development of D. *melanogaster* under low oxygen concentrations and for adult flies to recover from hypoxic exposure.

## Discussion

Oxygen is the final acceptor of the respiratory chain ensuring high yield of ATP production in eukaryotic cells. Upon oxygen reduction, cells adapt to lowered ATP levels by simultaneously reprogramming gene expression and reducing ATP demand. Gene reprogramming enables cells to modulate negatively or positively the expression of genes, several of which sustain the metabolic shift to elevated lactic glycolysis as an alternative ATP producing pathway. Concomitantly, major energy-consuming processes including protein synthesis are strongly dampened. It has been shown in human cells that a selected set of mRNAs remains associated to the translation machinery under hypoxia and escape translation inhibition^34^. D. *melanogaster* is particularly resistant to oxygen deprivation as it is physiologically exposed to variable oxygen concentrations and all cells of the organism are exposed to the ambient oxygen concentration, making this species the ideal study model to characterize molecular processes underlying cellular adaptation to oxygen variations. In the present study, we show that as in mammalian cells, protein synthesis is strongly reduced in *Drosophila* S2 cells due to a massive disassembly of polysomes. However, the mRNA encoding LDH is actively translated under those conditions, thereby contributing to the overall increased synthesis of LDH in hypoxic conditions. In terrestrial animals, *Ldh* plays an essential role under reduced oxygen concentration by contributing to the metabolic shift to lactic glycolysis. Its expression is strongly induced at the transcriptional level under hypoxia^35^. Here, we show that the 3’UTR of *Drosophila Ldh* mRNA is necessary and sufficient to activate translation under hypoxia. While 3’UTR-dependent translation under hypoxia of selected mRNAs has been previously reported in mammalian cells, our study points for the first time to the role of *Ldh* mRNA 3’UTR in translation activation. The study of Uniake et al. (2012)^14^ identified CGG motifs as translation-activating elements under hypoxia. Two copies of CGG trinucleotides are present in Drosophila *Ldh* mRNA 3’UTR. However, this motif is absent in mouse and human *Ldh* paralogs. The precise nature of the *cis*-regulatory element controlling the Drosophila *Ldh* mRNA translation remains to be established.

Our data identify the translation initiation factor eiF4EHP as an important element of the translation machinery under hypoxia. Hence, eiF4EHP deficiency strongly impairs Drosophila development under hypoxia and significantly reduces fly recovery after hypoxic treatment, thereby demonstrating the physiological importance of eiF4EHP in Drosophila adaptation to oxygen variations. eiF4EHP belongs to the family of Cap-binding eiF4E proteins. In contrast to the canonical cap-binding protein eiF4E1, eiF4EHP displays a reduced affinity to the cap structure and does not interact with eIF4G^36^. So far, it has been reported to act as a translation repressor by interacting with RNA-binding proteins recruited on mRNA 3’UTR and competing with eiF4E1 binding to the cap structure of these mRNA (see ^37^ and ^38^ for review). As eiF4EHP does not interact with eiF4G1 to form the active eiF4F cap-binding complex, the recruitment of 43S preinitiation is blocked. In contrast, eiF4EHP is weakly bound by 4E-BP proteins, thereby escaping from the 4E-BP-mediated sequestration induced by hypoxia^39^. Interestingly, the human homolog of eiF4EHP, eiF4E2, has been shown to be involved in translation in human cells under hypoxia^14^, indicating that the translation activating role of eiF4EHP in hypoxia has been conserved across evolution. In this context, human eiF4E2 can recruit eiF4G3 to promote hypoxic translation^40^. We observed that inhibition of Drosophila paralogs of eiF4G3 (eiF4G2, Nat1) do not impair translation under hypoxia (data not shown). Therefore, Drosophila eiF4EHP and mammalian eiF42 could recruit translational initiation complexes of different compositions.

Several eiF4EHP-binding proteins not related to eiF4G have been identified both in *Drosophila* and mammals. However, whether acting on global translation or regulating specific mRNA, most of these factors mediate translational repression and contribute to eiF4EHP migration in cytoplasmic granules such as P bodies or stress granules (reviewed in^38^). Our results showing eiF4EHP distribution in cytoplasmic granules distinct from P bodies and stress granules further indicate that eiF4HP plays a different role in the context of oxygen deprivation.

In human, the threonyl-tRNA synthetase (TRS) interacts with eiF4E2 to form a cap-dependent alternative translational initiation complex^15^. However, this interaction is vertebrate-specific and therefore cannot account for the translational activity of eiF4EHP described here.

The 3’UTR-dependent translation mechanism under hypoxia reported by Uniake et al.^14^ in human cells was shown to involve the formation of a complex composed of eiF4E2, eiF4A, HIF2alpha and the RNA-binding protein RBM4. The interaction of RBM4 with CGG motifs in mRNA 3’UTR contributes to selective mRNA translation by its ability to recruit the hypoxia-induced HIF2alpha and form of a translation-competent complex including eiF4E2. HIF2alpha belongs to the family of the HIF transcription factors activating the expression of several genes at the transcriptional level upon oxygen deprivation in mammals. It is worth noting that *Drosophila* has no homolog of HIF2alpha with the HIF1alpha protein Sima being a master regulator of hypoxia-induced genes^41^. While it is found to interact with several components of the translation machinery ^42^, Sima has not been identified to interact with eiF4EHP. Here, we show that *Ldh* mRNA is translated by a 3’UTR-dependent mechanism and involves the direct binding of eiF4EHP. At this point, we can hypothesize that eiF4EHP interacts with factor(s) promoting the recruitment of a translation initiation complex by a 3’UTR-specific mechanism. Further work is in progress to characterize this newly identified function of Drosophila eiF4EHP.

## Materials and methods

### Reagents

DNA oligonucleotides were purchased from Sigma-Aldrich. Puromycin, hygromycin, and neomycin were purchased from InvivoGen. pAC-sgRNA-cas9-puro and pAC-y1sgRNA-cas9-puro plasmids^43^ were obtained from Addgene (# 49330 and 49331). Anti-LDH (H-160) antibody was purchased from Santa Cruz (sc-33781). Anti-LDHA was purchased from Invitrogen (PA5-26531) or from Abcam (Ab130923). Anti-actin (A2066) and anti-puromycin antibodies (clone 12D10, Merck) were purchased from Sigma-Aldrich. 4EHP mouse monoclonal antibody was produced as previously described^44^. HRP-conjugated anti-mouse IgG and anti-rabbit IgG were purchased from Amersham Biosciences and Jackson ImmunoResearch Inc., respectively.

### Cell culture and hypoxia treatment

Non-adherent *D. melanogaster* S2 cells were originally purchased from Invitrogen and grown in Schneider’s Drosophila medium (Invitrogen) supplemented with 10% heat-inactivated fetal bovine serum under normoxia (21% O_2_) or hypoxia (1% O_2_) induced by N_2_ gas flow controlled by an oxygen sensor (CoyLab).

### Puromycin labeling

S2 cells (3.10^6^ in 2 ml of Schneider medium) were incubated at 21 or 1% O_2_ for 24h. Puromycin (5 μg/ml) was added to cultures and cells were further incubated for 20 min before lysis in EBC buffer (Tris 50mM pH8, NaCl 125mM, NP-40 0,5%, Na B glycerophosphate 50mM, NaF 50mM, Na_3_VO_4_ 200μM, protease inhibitor cocktail (Roche)). Equal amount of protein extracts was analysed by western blot analysis with anti-puromycin antibody. Actin was measured as loading control.

### Protein extraction and Western blot

Protein extracts were prepared by cell lysis in EBC buffer and protein concentration was measured with Bio-Rad protein assay dye reagent concentration. Proteins from sucrose gradient fractions were extracted by TCA precipitation. Western blots were performed according to standard techniques using the indicated antibodies. Signals were detected using the supersignal West pico detection kit (Thermofisher) and images were acquired and quantified with a Licor Odyssey FC imaging system.

### Generation of stable cell lines

The 3’UTR of *Ldh* gene was cloned into the pMT-luciferase plasmid^19^ and transfected in combination with a neomycin-resistant plasmid (1/20 ratio). Cells were selected for at least 3 weeks with neomycin (200 μg/ml). When indicated, cells were induced by CuSO_4_ (0.5mM) for 3h. ei4EHP and eIF4E6 Cas9 and CTRL KO S2 cell lines were generated by transfection of a pAC-Cas9-puro plasmid^43^ in which the *eiF4EHP* (GGAGCTCCCGGTAGGGCTTCAGG), *eiF4E6*(GGGGATCCGCCGAACAAGGG) or *Yellow* (Addgene #49331) guide sequences were inserted. Cells were selected with puromycin (5μg/ml) for at least 3 weeks. Subcloned cells were selected by limit dilution. All transfections were performed with Fugene HD according to the manufacturer’s instructions (Promega).

### Sucrose gradient polysome fractionation

Stably transfected S2 cells were cultured at 21% O_2_ (60. 10^6^ cells) or at 1% O_2_ (120.10^6^ cells) for 24 hours. Cells were treated with cycloheximide (10μg/ml) for 5 min., harvested and placed on ice for 5 min. Cell pellets were resuspended in 500 μl of polysome lysis buffer (Hepes 25mM, KCl 100mM, MgCl_2_ 5mM, Nonidet P-40 0.5%, heparin 2μg/ml, cycloheximide 10μg/ml, RNaseOUT (Invitrogen) 100u/ml), incubated on ice for 5 min. and centrifuged at 12000g for 5min. Lysates were loaded onto linear 15% to 50% sucrose gradients prepared in polysome lysis buffer and centrifuged at 39,000 rpm using a SW41 rotor for 2h at 4°C. Fractions were collected with an ISCO Density Gradient Fractionation System (Brandel).

Adult flies were kept in normoxia or exposed to 1 % O_2_ for 24h. Approximately 250 flies per conditions were lysed with a Dounce homogenizer (pestle B) in lysis buffer (Tris 10mM pH7.5, NaCl 150mM, MgCl_2_ 10mM, Igepal CA630 1%, Triton X100 1%, Na deoxycholate 0.5%, DTT 2mM, cycloheximide 200μg/ml, DNAse I 2u/ml, RNaseOUT 40u/ml, 1X Protease inhibitor cocktail EDTA free, heparin 10u/ml) and incubated on ice for 20min. Extracts were cleared by two rounds of centrifugation at 10000g for 5min. Supernatants were loaded on 15-50% linear sucrose gradients, ultracentrifuged at 39,000 rpm for 2h at 4°C. After fractionation, total proteins were recovered by TCA precipitation, rinsed with acetone, neutralized with Tris (1M) and resuspended in Laemmli buffer.

### RNA extraction and northern blot

Total RNA and RNA from sucrose gradient fractions were extracted by the Tri-reagent method. RNA from sucrose gradient fractions with heparin (55 u/ml) were extracted with the RNA Purification kit (Sigma RNB100-50RXN). Northern blots were performed as described elsewhere^45^. Briefly, blots were hybridized with RNA probes *in vitro* transcribed with ^32^P-labeled UTP (PerkinElmer Life Sciences). Radioactive signals were detected with a phosphorimager and signals were quantified with Quantity ONE software (Bio-Rad).

### RNA quantification by real-time PCR (qPCR)

cDNA was synthesized with 1 μg of total RNA using the Prime ScriptTM RT Reagent Kit (Takara) according to manufacturer’s instructions. Quantitative PCR was performed on a StepOnePlus real-time PCR system (Applied Biosystems) using SYBR® Premix Ex TaqTM II (Takara) according to manufacturer’s instructions. Expression levels were normalized to Rpl32 (ΔΔCT). The sequences of the primers used were Luc: 5′-gcctgaagtctctgattaagt-3′/ 5′-acacctgcgtcgaagatgt-3′; Ldh: 5′-caccgacatcctcaagaacat-3′/5′-gggattggacaccataagca-3′; Rpl32: 5′-gacgcttcaagggacagtatctg-3′/5′-aaacgcggttctgcatgag-3′; CAT: 5′-tccatgagcaaactgaaacgt-3′/5′-tgtgtagaaactgccggaaatc −3′.

### Reporter gene assays

Ldh and SV40 3′UTR sequences were subcloned into a vector expressing the luciferase coding sequence under control of the Ldh promoter (see figure legends). Cells were transiently transfected with these reporter constructs in combination with a Renilla luciferase control plasmid. Twenty-four hours after transfection, cells were exposed to variable O_2_ concentrations for another 24 hours. Cells were lysed and luciferase activity was measured using the Dual-Luciferase Reporter Assay (Promega) according to manufacturer’s instructions.

### Immunofluorescence and FISH experiments

For immunofluorescence experiments, S2 cells (10^6^ in 1 ml) were seeded on microscope slides previously treated with concanavalin A (0.5 μg/ml) for 15 min. After the indicated treatments, cells were fixed and permeabilized as previously described^44^. After antibody staining, slides were mounted in fluorescent mounting medium (DAKO, Glostrup, Denmark) supplemented with 100 pg/ml 4’,6’-diamidino-2-phenylindole (DAPI) and sealed with nail polish. For co-localization experiments, cells were transfected with pMT-GFP-Rox8 or pMT-GFP-Dcp1 using Fugene HD according to the manufacturer’s instructions and treated overnight with CuSO_4_ (0.5mM) before plating on slides.

For polyA^+^ RNA FISH experiments, cells were permeabilized with cold methanol after fixation and were subsequently washed 3X with 2XSSC and hybridized with biotinylated oligodT (1ng/μl) in hybridization buffer (final composition: dextran 10%, BSA 0.5%, E. coli tRNA 1mg/ml, 2XSSC, Formamide 25%) overnight at 42°C. After wash (3×) with 2XSCC, slides were incubated with AlexaFluor 594-coupled streptavidin (Thermofisher) at 1/1000 dilution for 1h at room temperature. Signal amplification was performed by incubation with biotin-coupled anti-streptavidin (1/1000) in 2xSSC-Tween20 0.2% followed by incubation with AlexaFluor 594-coupled streptavidin. Slides were washed and mounted in DAKO supplemented with DAPI.

FISH/IF experiments were performed according to the same procedure combining the amplification steps with protein staining in 2xSSC-Tween20 0.2% followed by incubation with AlexaFluor594-coupled streptavidin and AlexaFluor488-coupled donkey anti-mouse IgG (Thermofisher) (1/1000) in 4xSSC-Tween 20 0.2%.

### Image acquisition and processing

Images were acquired with a Zeiss LSM710 confocal microscope equipped with a 63×/1.4 oil-immersion Plan-Apochromat objective. Maximum intensity projection images of Z-stacks were generated with Zeiss Zen Blue software. Signal analysis and quantification was performed with ImageJ software. Signal intensity was calculated as the signal mean intensity in individual cells subtracted from signal background. Signal heterogeneity was assessed by calculating the standard deviation of signal intensity in individual cells divided by the square root of the signal intensity. Co-localization of signals was analysed using RGB Intensity Profile tool in ImageJ on manually drawn lines across imaged cells.

### CLIP experiments

Cells cultured in normoxia (21% O_2_) or hypoxia (1% O_2_, 24h) were washed once with 1 ml of cold PBS and plated on a new plate before UV crosslinking (254 nm, 150 mJ/cm^2^) in a UV Stratalinker 1800 (Stratagene). The cells were collected and lysed in 0.5ml of lysis buffer (Tris-HCl 50mM pH 7.5, EDTA 0.5mM pH 8, NP-40 0.5%, glycerol 10%, NaCl 120mM, sodium b-glycerophosphate 50mM, Na_3_VO_4_ 200μM, NaF 50mM, protease inhibitor (50× diluted), Ribolock (4000× diluted)), incubated on ice for 10 min. and centrifuged at 13.000 rpm for 10 min. at 4°C. 50 μl were kept as input and the rest was incubated overnight with 50 μl of protein G Sepharose beads coated with anti-eiF4EHP antibody at 4°C. Beads were washed twice with NT2 buffer (Tris 50mM pH 7.4, NaCl 150mM, MgCl_2_ 1mM, NP-40 0.05%) before RNA release by proteinase K (0.6mg/ml) treatment for 30 min. at 37°C and 3 min. at 95°C. RNA was extracted with 0.5ml of Trizol spiked with diluted *in vitro* transcribed CAT mRNA. Total RNA was resuspended in 30μl of water. 6μl of each sample were treated with 1μl of DNase I (Thermo Scientific) for 30 min. at 37°C before inactivation in EDTA (5mM, pH8) for 10min. at 65°C. Reverse transcription and qPCR analysis were performed as described above.

### Fly strains and maintenance

The fly strains used in the experiments were obtained from the Bloomington Drosophila Stock Center (BDSC), including: Canton-S (RRID: BDSC_64349); y^1^ v^1^ (RRID: BDSC_1509); y^1^ sc* v^1^ sev^21^; P{y^+t7.7^ v^+t1.8^=TRiP.GL01035}attP2 (RRID: BDSC_36876, named as “4EHP RNAi”); P{w[+mC]=tubP-GAL4}LL7/TM3, Sb^1^ Ser^1^(RRID: BDSC_5138 named as “Tub-Gal4”); y[1] sc[*] v[1] sev[21]; P{y[+t7.7] v[+t1.8]=VALIUM20-mCherry}attP2;(RRID:BDSC_35785 named as mCherry_RNAi). All the *in vivo* experiments were carried out at 25°C under indicated O_2_ concentrations.

### Negative geotaxis assay

Age-matched controls and *eiF4EHP* knock-down female flies were exposed to 1% O_2_ environment for 24h and returned to 21% O_2_. Measurement of fly mobility was performed after 40 min. and 48h of recovery time on 20 individuals per condition as previously described^46^.

### Statistics and graphs

Graphs were made using GraphPad Prism 8. Other statistical tests were performed as indicated in the text.

## Acknowledgements

We thank Dr Elsa Lauwers for help and advice with Drosophila experiments. Manfei Liang was supported by the Chinese Research Council, the Fonds van Buuren, and the Fonds Rose et Jean Hoguet, Clara Hody was funded by the FNRS (aspirant grant). This work was supported by a grant from the European Regional Development Fund (FEDER), the Belgian Fonds de la Recherche (FNRS, PDR 23605960 and CDR J.0079.21F), the Fonds Jean Brachet and the International Brachet Stiftung.

## Author contributions

M.L, C.H, V.Y, R. S, Y.S, X.L, X.T, R.M, M.P, F.A, and L.C carried out the experiments. C.G and V.K wrote the manuscript with support from M.L, V.Y and C.H. C.G and V.K supervised the project.

